# Unity and diversity in working memory load: Evidence for the separability of the executive functions updating and inhibition using machine learning

**DOI:** 10.1101/389395

**Authors:** Tanja Krumpe, Christian Scharinger, Wolfgang Rosenstiel, Peter Gerjets, Martin Spüler

## Abstract

**Abstract:** *Objective:* According to current theoretical models of working memory (WM), executive functions (EFs) like updating, inhibition and shifting play an important role in WM functioning. The models state that EFs highly correlate with each other but also have some individual variance which makes them separable processes. Since this theory has mostly been substantiated with behavioral data like reaction time and the ability to execute a task correctly, the aim of this paper is to find evidence for diversity (unique properties) of the EFs updating and inhibition in neural correlates of EEG data by means of using brain-computer interface (BCI) methods as a research tool. To highlight the benefit of this approach we compare this new methodology to classical analysis approaches.

*Methods:* An existing study has been reinvestigated by applying neurophysiological analysis in combination with support vector machine (SVM) classification on recorded electroenzephalography (EEG) data to determine the separability and variety of the two EFs updating and inhibition on a single trial basis.

*Results:* The SVM weights reveal a set of distinct features as well as a set of shared features for the two EFs updating and inhibition in the theta and the alpha band power.

*Significance:* In this paper we find evidence that correlates for unity and diversity of EFs can be found in neurophysiological data. Machine learning approaches reveal shared but also distinct properties for the EFs. This study shows that using methods from brain-computer interface (BCI) research, like machine learning, as a tool for the validation of psychological models and theoretical constructs is a new approach that is highly versatile and could lead to many new insights.

## 1. Introduction

Working memory (WM) is a core construct in cognitive psychology that can be represented as a set of functions, which are responsible for the temporary storage and the simultaneous processing of information during cognitive tasks (Miyake and Shah, 1999). The still dominant multicomponent model of WM by Baddeley and colleagues (2012; 1974) distinguishes three main components of WM, two of which are memory-related storage components for verbal and pictorial information. The third component is an attention-related central-executive control structure, which comprises different executive functions (EFs) for coordinating the storage and transformation of information in memory cf., (Baddeley, 1996, 2007). EFs play a pivotal role in many more recent WM theories e.g., (e.g., Barrouillet et al., 2004; Cowan et al., 2005; Kane et al., 2004; Oberauer, 2009; Unsworth and Engle, 2007) and (Cowan et al., 2014). According to these theories, the attentional control processes required to pursue WM tasks are responsible for the severe limitation of human WM. Accordingly, individual differences in WM capacity are traced back to individual differences with regard to these EFs (Kane et al., 2004; Unsworth and Engle, 2007).

Based on a latent-variable analysis of a bundle of executive tasks, Miyake et al. (2000) developed one of the best-known models describing the structure of different EFs. According to this model, the three core EFs are named *updating, inhibition* and *shifting*. Updating is described as a process of keeping information in WM up to date for a certain period of time (Ecker et al., 2010), for instance by manipulating this information or by loading new information and unloading older pieces of information. Shifting can be described as the adjustment of task rules currently kept in mind to new circumstances (Monsell, 2003). Inhibition refers to preventing (currently irrelevant) information or action tendencies from getting access to WM while being involved in a cognitive task (Diamond, 2013). Miyake and colleagues (Miyake and Friedman, 2012; Miyake et al., 2000) describe the relation between the three core functions in terms of unity and diversity, as they share many properties but also have distinct characteristics as individual functions.

### 1.1. Previous work: Neurophysiological assessment of WML/EFs

The assessment of working memory load (WML) in a global manner, without differentiating neural correlates with respect to EFs, has been done in many studies before. The amount of load can be measured and quantified in EEG data by means of neural correlates in the power spectra but also in behavioral data like reaction time and the ability to execute a task correctly. Neural correlates of WML in EEG data can be described especially by differences in the theta and alpha band power. During an increase of WML theta power has been reported to increase at frontal electrode sites, also known as event related synchronization (ERS) (Gevins et al., 1997; Jensen and Tesche, 2002; Missonnier et al., 2006). In contrast to that alpha power has been observed to decrease at paretial electrode sites during an increase in WML, which is also known as event related desynchronization (ERD) (Gevins et al., 1997; Krause et al., 2010; Stipacek et al., 2003). With regard to pupil diameter, it is known that an increased level of WML is reflected in a larger diameter (Beatty and Lucero-Wagoner, 2000; Ewing and Fairclough, 2010a; Karatekin et al., 2007; Laeng et al., 2011). Regarding ERPs, a decrease in the P300 amplitude at parietal sites for increased WML has been reported (Allison and Polich, 2008; Brouwer et al., 2012; Pratt et al., 2011; Watter et al., 2001). Especially spectral features have been used so far, to categorize the current amount of load into high and low e.g., (Brouwer et al., 2012; Putze et al., 2010; Walter et al., 2017), by means of machine learning, thus in an automated manner, which has been shown to work reliable and successfully.

The unity and diversity of EFs as individual components of WM, describing shared and unique properties of the EFs, has so far mainly been investigated by means of statistical analyses of behavioral data in either healthy subjects or patients with frontal lobe impairments, see e.g., (Burgess, 1997; Shallice and Burgess, 1993) and (Shallice and Burgess, 1991). So far, it remains unclear whether the shared and unshared variances of performance in different EF tasks can be mapped either, on a common attentional and limited resource in the brain or, on EF-specific brain functions. One example of studies that aimed to answer this question by Collette et al. (2006; 2005) explored neural substrates for the EFs updating, inhibition and shifting by means of positron emission tomography (PET). Via conjunction and interaction analysis they compare a battery of tasks for each EF to reveal both unity and diversity aspects within the collected brain data. An assessment of differences in neural correlates by means of EEG data, that allow a within subject and not only between subject comparison between EFs has, to the best of our knowledge, not been done before, except for one study by Scharinger et al. (2015).

Scharinger et al. (2015) integrated an n-back task (imposing updating demands) within a flanker task (imposing inhibition demands) to study the relation between the two EFs updating and inhibition simultaneously within one perceptually and motorically highly controlled task. In their study they manipulated demands on the two EFs independently and analyzed indicators of WML, both at the behavioral level (reaction times (RTs) and accuracies) and at the physiological level (pupil diameter, event-related potentials in the EEG (ERPs) and EEG power spectra) to develop a detailed and brain-related account of how load on different executive functions is interrelated.

### 1.2. Foundation: Study Scharinger and colleagues

The above mentioned study by Scharinger et al. (2015) presented a new study design which seems very promising with respect to disentangle individual properties of EFs, in this specific case updating and inhibition. Demands on both EFs are induced and manipulated within the same experimental design by using an n-back task with a simultaneously presented flanker. The design ensures that visual presentation and the motoric requirements for the subjects are kept steady and simple, reducing the non-EF variance in the data to a minimum. The n-back task is a widely used WM task in neuropsychological research for studying effects of WM updating, e.g., (Chatham et al., 2011; Chen et al., 2007; Ewing and Fairclough, 2010b; Gevins and Smith, 2000; Jonides et al., 1997; Krause et al., 2000; Owen et al., 2005; Pesonen et al., 2007). The simultaneously used flanker task is known to induce inhibition demands, when the flanking letters are incongruent (i.e different) to the central and task relevant letter. The perceptual differences lead to interference effects (Eriksen, 1995; Sanders and Lamers, 2002) which, in turn, provokes additional executive control to overcome the interference.

Scharinger et al. (2015) showed, first, that all load-related measures yielded the expected outcomes for an increase of the n-back levels (load on working memory updating) confirming the sensitivity of the used indicators as measures of WM updating load. Second, all measures were also sensitive to inhibitory demands. Thus the task design has been proven to be legitimate. The most important finding by Scharinger et al. (2015) was, however, that the flanker interference effect did decrease under high WM updating load as shown by the outcomes of all measures. Only for low load on updating the flanker interference effect led to increased RTs and increased pupil dilation as well as to decreased upper alpha frequency band power and decreased P300 mean amplitude. In contrast, under high load on updating, no significant flanker interference effect was observed (RT, pupil dilation, P300) or the effect even was reversed (EEG upper alpha frequency band power). Scharinger et al. (2015) hypothesized that if the different executive functions proposed by Miyake et al. (2000) were rather separate functions, additional load on inhibition due to a flanker conflict within the n-back task paradigm should lead to simple additive effects on all n-back levels for all of the aforementioned measures. In contrast, if these three executive functions were more closely intertwined, i. e., rely on a common attentional resource, it would be expected to observe interaction effects in the load related measures for additional load on inhibition. Thus, the study results suggest, that the EFs updating and inhibition might share underlying network structures that serve controlled attention (unity of executive functions).

### 1.3. Aim of this study

The aim of this study is to evaluate if the EEG data provided by Scharinger and colleagues also reveal evidence for the diversity of the EFs updating and inhibition. Generally, diversities in terms of neural correlates in EEG data are evaluated by calculating an averaged form of all relevant trials, since noise, random effects and inter subject variability is reduced and only the characteristic properties remain. In special cases in which individual differences are of importance to define a certain mental state, the possibility for a distinction between states, could get lost though with this classical analysis approaches. Using machine learning approaches to separate mental states individually for each subject on the basis of their EEG signals might be a solution for this problem. We want to act on this suggestion and aim to find out if the two EFs are distinguishable by means of machine learning approaches. In addition, a method for proper feature interpretation will be applied to the machine learning approach to reveal which features are of importance in the distinction of the EFs. Thus, revealing which features can be correlated with diversity. The results will be compared to conventional EEG analysis techniques to identify the advantages of the proposed analysis methods.

## 2. Material and Methods

### 2.1. N-back-flanker Task design

In this paper, we will reanalyze physiological data from an experiment reported by Scharinger et al. (2015) that uses the integrated n-back flanker task to study interactions between the two EFs updating and inhibition. The stimuli consisted of the four letters, S, H, C and F. For each trial, one out of these four letters was randomly chosen and presented centrally on the screen either flanked by the same letters (congruent condition without inhibition demands, the same letter appeared three times both on the left and right sides of the centered target letter, e.g., HHH H HHH) or by randomly chosen different letters (incongruent condition with inhibition demands, one of the three remaining letters appeared three times both on the left and right sides of the centered target letter, e.g., FFF H FFF). All letters were presented in gray on black backgrounds in Arial at 25-point font size. Each stimulus was shown for 500 ms, followed by a black screen for 1500 ms. Thus, each trial lasted 2000 ms. For a schematical overview see Figure 1. In the experiment, three levels of updating demands were implemented (n = 0, 1, and 2) in a block design. For each trial in a block the subjects indicated via key press (yes/no key) whether the central letter of the current trial was identical to (target) or was different from (nontarget) the central letter they had seen in the sequence n steps back. In the 0-back condition, before the stimulus sequence started a randomly chosen letter (S, H, C, or F) was displayed as the n-back target letter for the whole block (no updating required). During the following stimulus sequence, each time this letter occurred as the central letter, subjects had to press the yes key, in all other cases, the no key. Answers and reaction times of the subjects were recorded.

**Figure 1.**
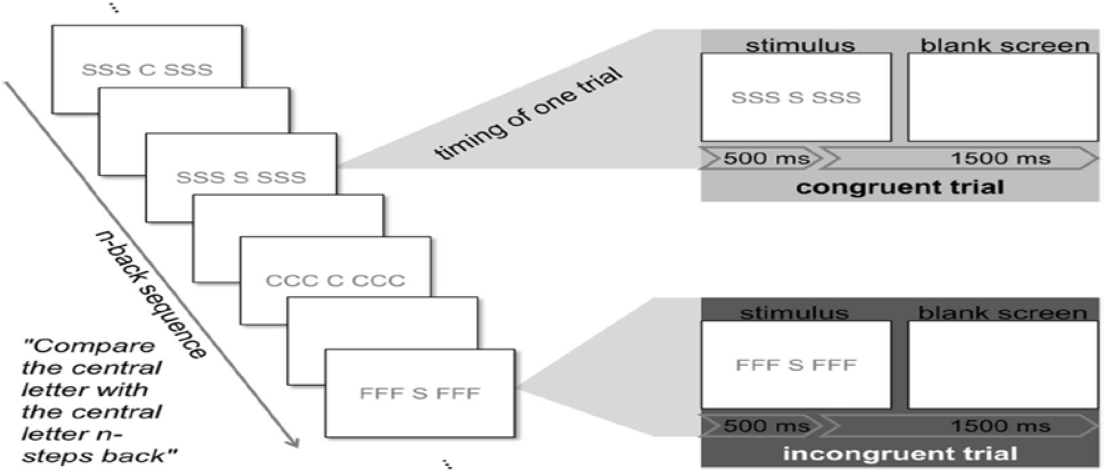
Experimental design and flow from Scharinger et al. (2015). An n-back sequence was presented in blocks, accompanied by flanker letters to the right and left of the n-back stimulus. All trials lasted two seconds of which 500 ms were stimulus presentation and 1500 ms a blank screen before the next stimulus appeared.

The n-back stimulus sequences were presented in blocks. In accordance with the traditional n-back task design, each block of 154 trials consisted of one n-back level. Each n-back level was presented twice. Thus, subjects performed a total of six blocks. The sequence of blocks was randomly assigned for each subject, with the constraint that each n-back level was presented once before an n-back level was presented for the second time. One block lasted about 5 min. Within each block, half of the trials were targets, half of the trials were nontargets. About one third of the stimuli of each response category were incongruent (e.g., FFF H FFF), two thirds were congruent (e.g., FFF F FFF). The first four trials of each block were always congruent nontargets and were excluded from any further analyses. Stimuli were presented using E-Prime presentation software (E-Prime 2 Professional, Psychology Software Tools, Inc.) with predefined stimuli lists. The trial sequences within the blocks were pseudorandomized, to avoid attenuation of the interference effect for incongruent stimuli due to conflict adaptation processes (i.e., the so-called Gratton effect (Botvinick et al., 2001; Davelaar, 2012; Gratton et al., 1992)), incongruent-incongruent stimuli sequences were excluded in advance during construction of the stimuli lists. To further avoid any Gratton-like effects, congruent trials following incongruent trials were excluded from any further data analyses. In addition to that, in each block 20 stimuli consisting of 10 targets and non-targets were randomly chosen and replaced by stimuli without a central letter (i.e., 10 targets and 10 nontargets per block consisted only of the flankers on both sides of a gap). In these cases, we instructed the subjects to rememberthe flanker letters of the current trial for the following comparison. By means of this instruction, we wanted to avoid subjects becoming increasingly unaware of the flanker stimuli during the course of a block. We excluded these gap stimuli and the two immediately following stimuli from any further analyses. At the beginning of the study, subjects performed training blocks for each n-back level. Training was repeated until subjects reached an accuracy of at least 60 percent correct responses. During training, subjects accuracy was displayed at the end of a block to give them feedback regarding their performance. No feedback was given during the actual task presentation.

### 2.2. Data recording

22 subjects (21 right handed, 12 females) participated in the study, for which they were reimbursed with 8 € per hour. All subjects had normal or corrected to normal vision and no reported neurological disorders. The study was approved by the local ethics committee and written informed consent was obtained from all participants. 32 electrodes (Acticap Brain Products) were used for the recording at a sampling rate of 500 Hz and placed according to the international 10/20 system (Jasper, 1958) with the reference at right mastoid and the ground electrode at AFZ. In addition to EEG recordings an eyetracking system (SMI iView X 2.7.13) was used to record subjects’ pupil diameter with a sampling rate of 250 Hz. For further details considering the technical setup see Scharinger et al. (2015).

### 2.3. Preprocessing of the data

For the physiological data, only artifact-free trials with correct responses were used for data analysis, with an additional exclusion of trials that might be effected by any Gratton-like effect (see Section 2.1 above). The data was bandpass filtered between 0.4-40 Hz and re-referenced to common average. To remove artifacts a threshold of 100 *μV* was chosen and all trials exceeding this level were discarded. Trials including eye movement artifacts were corrected using independent component analysis (ICA) (rejection by visual inspection). In addition, a baseline correction using a −250-0 ms prestimulus section was performed on all trials. These preprocessing steps resulted the following amount of trials per factor level: 0-back, congruent, M = 54.05, SD = 5.24; 0- back, incongruent, M = 65.68, SD = 7.08; 1-back, congruent, M = 52.77, SD = 6.68; 1- back, incongruent, M = 64.54, SD = 9.50; 2-back, congruent, M = 44.05, SD = 8.89; 2- back, incongruent, M = 55.05, SD = 11.07.

### 2.4. Neurophysiological analysis

Scharinger et al. (2015) conducted 2×3-factorial ANOVAs to test for interaction effects when combining WM updating load with flanker interference. These ANOVAs demonstrated that flanker interference effects decreased under high WM updating load, indicating a close connection of the two EFs in terms of their physiological correlates. In addition to these findings, the current reanalysis will test whether there is also evidence in the data for distinct physiological correlates indicating the diversity of the two EFs updating and inhibition. Updating demands are supposed to be induced when the n-back level is greater than 0 and inhibition demands are supposed to be induced when the flanker is incongruent to the n-back stimulus. In our reanalysis, a machine learning approach based on SVM classification will be used to separate load on these two EFs. In particular, it will be tested how good individual trials can be classified into one out of four factor levels based on the physiological responses collected during an epoch of 0-1000 ms after stimulus onset:

- Baseline trials (BL) without load on EFs (0-back, congruent)
- Inhibition trials (Inh) with load on inhibition only (0-back, incongruent)
- Easy updating trials (Up1) with load on updating only (1-back, congruent)
- Difficult updating trials (Up2) with load on updating only (2-back, congruent)

The remaining two factor levels containing mixed trials that impose both types of load on EFs simultaneously (i.e., 1-back and 2-back with incongruent flanker) will not be analyzed by this classifier approach as they do not represent clear categories with regard to the type of load they impose on EFs. Three types of physiological features were used for classification: ERPs, EEG power spectra, and pupil diameter. For the investigation of ERPs the grand average over all subjects and trials was computed for each of the four factor levels separately. Based on the results of Scharinger et al. (2015), the ERPs at electrode positions FZ, CZ, PZ were of major interest in the context of this task as they yielded effects with regard to frontal positivity and also with regard to the amplitude of P300 potentials. Apart from the ERP analysis, the data was also analyzed in the frequency domain to get insights into the spectral properties of the two executive functions. For the calculation of the power spectra Burgs maximum entropy method was used with a model order of 32 and a bin size of 1. With regard to the pupil diameter, the mean diameter for each trial during an epoch of 0-1000 ms was analyzed. To test whether the differences in the ERPs, power spectra, and pupil diameter between the factor levels are statistically significant a Wilcoxon ranksum test was conducted over all subjects and trials. The resulting p-values were Bonferroni corrected and the significance level was set to p < .05

### 2.5. Classification

To separate the EFs by means of machine learning, support vector machine (SVM) classification was chosen as a possible way to achieve this goal. A SVM with a linear kernel (C = 1) (Platt, 2000; Vapnik and Chervonenkis, 1974) was applied to differentiate between the four different factor levels introduced above (BL, Inh, Up1, Up2) using the libsvm implementation for Matlab (Chang and Lin, 2011; MATLAB, 2015). The classification between factor levels was conducted for the following pairs for each subject individually: Inh vs BL, Up1 vs BL, Up2 vs BL, Inh vs Up1 and Inh vs Up2. To ensure stable results a 10-fold cross validation for each comparison was implemented, dividing the available data into ten different test and training sets. The datasets (training set as well as the test set) were balanced for each comparison, by removing all spare trials if one of the classes had more trials than the other. Classification was performed on single trial basis for each comparison. For classification, three different types of features were used: ERPs in the time-domain, power spectra of the EEG data, and pupil diameter. The time domain features are based on 0-1000 ms epochs, starting at stimulus onset of the 17 channels (FP1, FP2, F3, FZ, F4, FC1, FC2, C3, CZ, C4, CP1, CP2, P3, PZ, P4, O1, O2). All other electrodes were discarded to reduce the influence of noise and artifacts in the data. Due to the sampling rate of 500 Hz one trial of ERP data is represented by 500 x 17 features. As a way to improve signal-to-noise ratio of the data, a spatial filtering method based on canonical correlation analysis (CCA) was applied (Spüler et al., 2014) with a filter size of 27×17. The filter aims to minimize the variance within a class and to maximize the variance between classes to improve the separability. Classification using features from the frequency domain was conducted on the power spectra between 4-13 Hz calculated on the same time frame (0-1000 ms after stimulus onset) with Burgs maximum entropy method for the same 17 channels (10 x 17 features). A third feature set included the pupil diameter of both eyes (0-1000 ms after stimulus onset, 500 x 2 features).

#### 2.5.1. Cross-class classification

Additionally, a cross-class classification analysis was conducted to extract information about the potential overlap of relevant feature characteristics between classes. Therefore, a classifier was trained on the distinction between executive function (EF1) and the baseline condition (BL) and tested on the other EF (EF2). If the relevant features and feature characteristics that distinguish EF1 and EF2 from BL overlap to a certain extend cross-classification accuracies should be significantly above chance level. Otherwise chance level accuracy is to be expected, as both available choices of labels are not applicable on the test trials.

#### 2.5.2. Neurophysiological interpretable features

To inspect the features used for the distinction, a method developed by Haufe et al. (2014) was used that transforms the weights of the SVM classifier into neurophysiological interpretable values. This transformation is a necessary processing step since multivariate methods like SVMs combine information from several channels to improve the signal to noise ratio, thereby preventing the possibility to directly interpret the involved parameters that lead to the decision of the classifier. The step that is done in a SVM is described by the authors as a backward model that transforms data *x*(*n*) to the optimized and separable form *s*(*n*) by multiplying a transformation matrix on the data (Eq. 1).

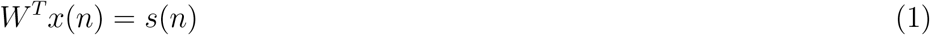

The transformation matrix represents the weights of the SVM which are mathematically optimized but cannot be interpreted in terms of the neurophysiological importance of the features that are used for the distinction of classes. To reveal the individual importance, the so called activation pattern *A* is calculated by multiplying the covariance matrices of the data with the weights of the SVM (see Eq. 2).

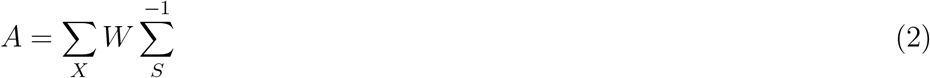

By calculating activation patterns the underlying neurophysiological patterns that are responsible for the distinction can be inspected, which can provide valuable information analyzing the unity and diversity of different EFs.

Statistical testing was performed on classifications for which the accuracies were close to chance level. The statistical significance of the results was determined by permutation tests with 1000 iterations (Good, 2013; Nichols and Holmes, 2002). The classification performance achieved in the permutations establishes an empirical null distribution on random observations, which can be used to determine significance boundaries. Therefore, in each iteration classification was performed in a 10-fold cross validation, but with randomly assigned class labels in the training set instead of the correct class labels. The achieved accuracy values were compared with the ones determined in the standard 10-fold cross validation in an ANOVA. Significance level was determined to be at p < .05, stating that the original classification performance is significant when the performance values are higher than the 95th percentile of the calculated empirical distribution.

## 3. Results

### 3.1. Neurophysiological analysis of the data

#### 3.1.1. P300 evaluation

ERPs were evaluated in averaged form for each condition at the electrode positions FZ, CZ and PZ. In Figure 2 the three subplots A, B and C show the grand average over all subjects for all four described factor levels (BL, Inh, Up1 and Up2). It can be seen that the conditions differ in amplitude, but the waveform remains rather constant. At position PZ (Figure 2 C) the waveform can be identified as a P300. Despite the differences within and across electrodes, the tendency: BL > Inh > Up1 > Up2, in terms of the amplitude can be observed at all electrode positions (with minor exceptions). Figure 2 also shows the time segments that differ significantly (p < .05) for each comparison between conditions at the three electrode positions analyzed. At position FZ almost no differences in the ERPs can be found across conditions, but at CZ and PZ significant differences in amplitude can be found across all comparisons between 200 and 750 ms.

**Figure 2.**
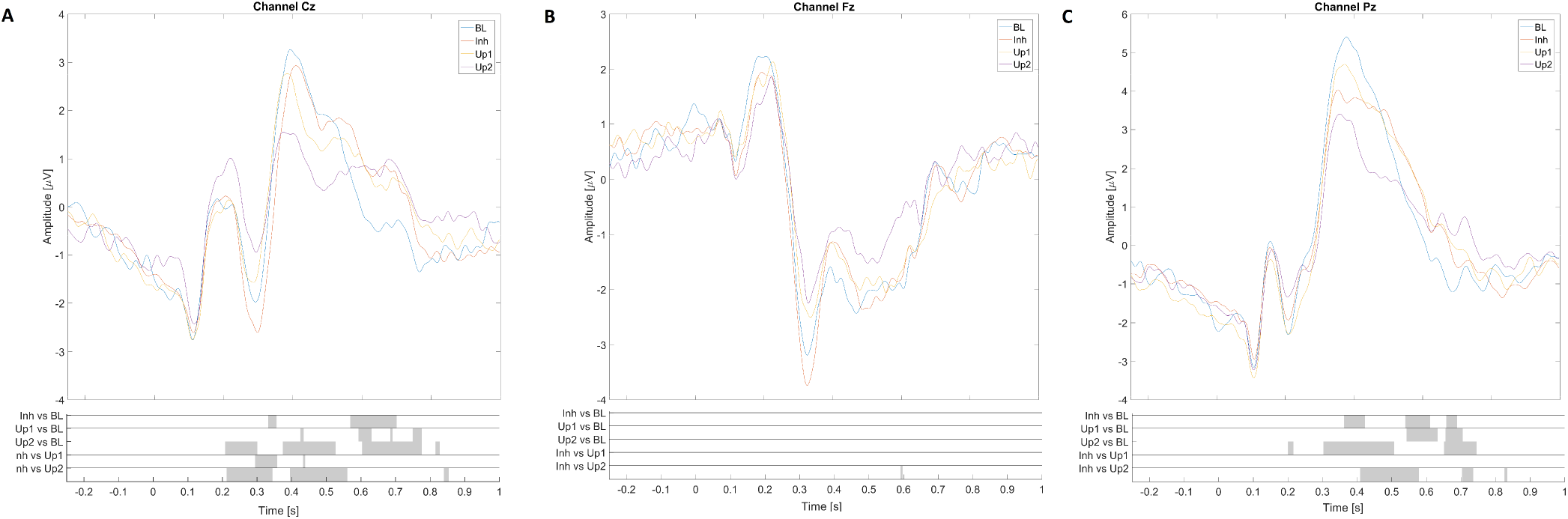
ERPs, 0-1000 ms from stimulus onset are depicted from the four different conditions BL (0-back, congruent), Inh (0-back, incongruent), Up1 (1-back, congruent) and Up2 (2-back, congruent) at positions Cz(A), Fz(B), Pz(C). The values represent the grand average over all subjects and trials. On the x-axis the time is represented in seconds, whereas the y-axis displays the amplitude of the ERPs in μV. Below the ERPs, the statistical evaluation is represented, revealing which time segments of the ERPs differ significantly between the conditions. Each grey bar states that the amplitude differs significantly (p < .05) for the respective comparison in the marked points in time.

#### 3.1.2. Alpha and theta power

Figure 4 shows the power between 4-13 Hz at different electrode positions calculated on a one second time frame after stimulus onset. Again the three positions FZ, CZ and PZ were chosen and can be found in this order in Figure 4 A, B and C. Frontal theta activity is visible at electrode position FZ as clear peak at 6 Hz (A), as well as parietal alpha activity at electrode position PZ (C) peaking at 11 Hz. The theta activity can be identified as an ERS since the power increases with the amount of induced load, whereas the alpha activity can be identified as an ERD as the power decreases with an increasing amount of load. Position CZ shows no notable spectral changes. Again the same pattern in value strength can be observed as in the ERPs and the pupil diameters. The higher the WML the higher and more concise the neural activity, in this case represented by the ERD/ERS in alpha and theta power. Frontal theta is highest for Up2 and lowest for BL, whereas Up2 shows the lowest parietal alpha power and BL the strongest. Figure 4 also shows the significant differences between the conditions for the three electrode positions between 4-13 Hz. Some comparisons show a lack of differentiable information in the power spectrum, but overall a broad number of features indicate that the EFs might differ from each other with regard to relevant features and feature characteristics. When comparing Inh vs Up1, Inh vs BL and Up1 vs BL not always distinct features can be found describing the variance of the EFs.

#### 3.1.3. Pupil diameter

The grand average of the pupil diameter (over all trials and subjects for 0-1000 ms after stimulus onset) can be seen in Figure 3 A. It reveals the same tendency as the ERPs for the individual conditions but inverted (increased amplitude of pupil diameter and decreased P300 amplitude with increasing load levels). Figure 3 B shows the variance of the pupil diameter values exemplary for the timepoint 200 ms. As the diameter seems to behave the same throughout all conditions (concerning the displayed waveform) this timepoint was chosen as a showcase to get an insight on how stable the pupil diameter values are. The variance seems to be quite high, but the tendency remains clearly visible.

**Figure 3.**
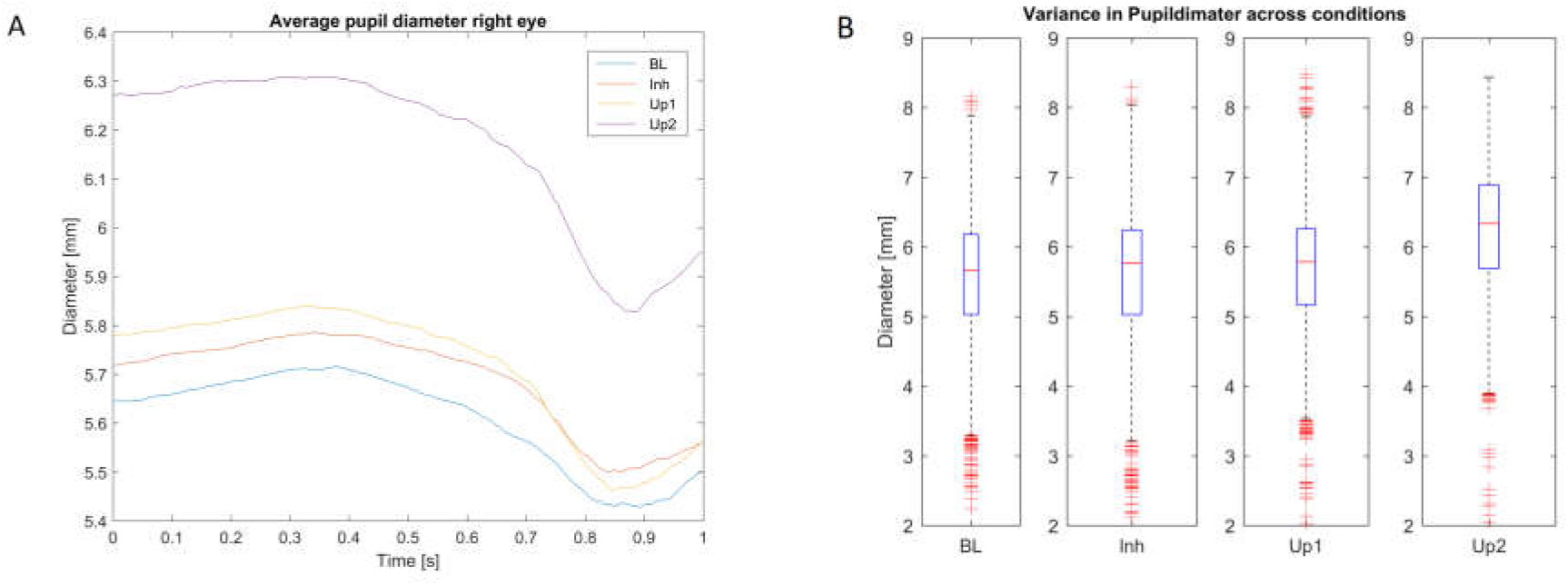
A: Pupil diameter of the left eye for the time frame of 0-1000 ms from stimulus onset sorted and averaged over all trials and subjects belonging to one condition. Displayed are the four different conditions BL (0-back, congruent), Inh (0-back, incongruent), Up1 (1-back, congruent) and Up2 (2-back, congruent). Each line displays the averaged diameter of one condition. B: The variance in the pupil diameter across all trials and subjects is represented in boxplots. Each box displays one condition at timepoint 0.2 s after stimulus onset. The bottom and top of the box represent the first and third quartiles, and the red band inside the box represents the median (the second quartile).

**Figure 4.**
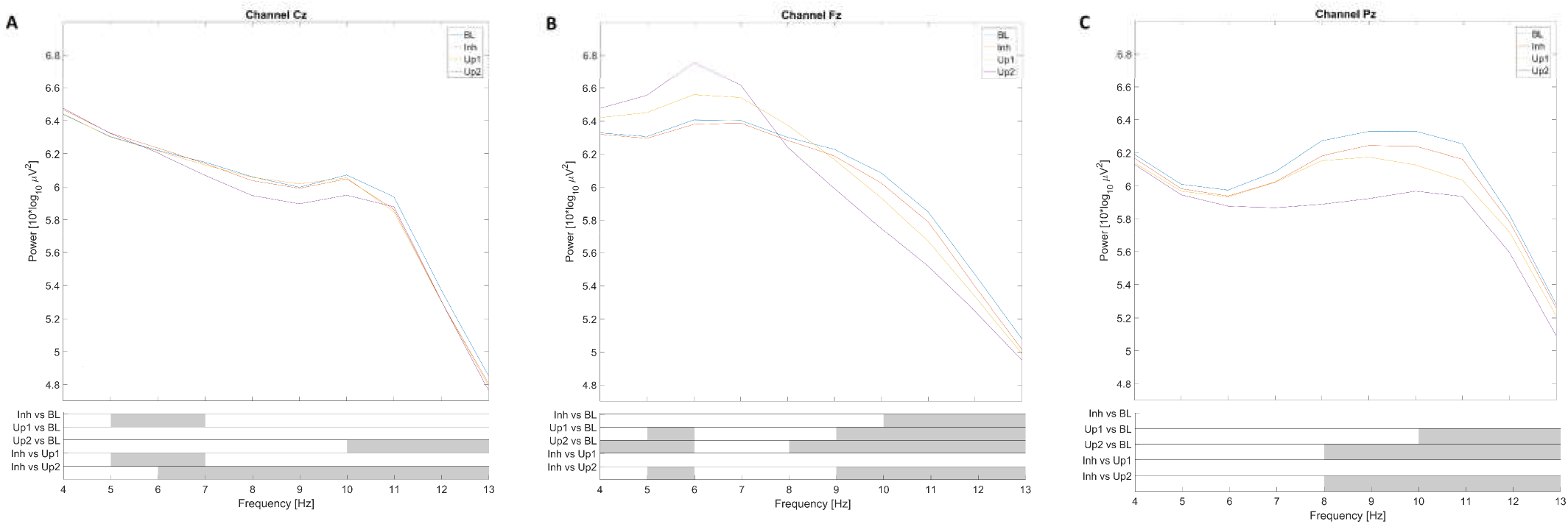
Power spectra of the four different conditions, BL (0-back, congruent), Inh (0-back, incongruent), Up1 (1-back, congruent) and Up2 (2-back, congruent) calculated from 4-13 Hz on the time frame 0-1000 ms from stimulus onset at positions Cz(A), Fz(B), Pz(C). Each line represents the grand average over all subjects and trials. Below the averaged power spectra, the gray bars represent the sections in the power spectra which differ significantly (p < .05) between the conditions.

### 3.2. Classification results

The feasibility to distinguish the three conditions imposing demands on EFs (Inh, Up, Up2) from baseline (BL) demands, as well as from each other, was evaluated using different classification approaches. In the following the accuracies of different choices of features will be reported, starting with ERPs, continuing with the pupil diameter of both eyes and the power spectrum between 4-13 Hz.

#### 3.2.1. Classification on ERPs

Classification was performed on the ERPs using the 17 mentioned channels. See Table 1 for the accuracy values that have been achieved in the 10-fold cross-validation, for each distinction. It can be seen, that the Up2 condition can be much better differentiated from BL and Inh demands than the Up1 condition. Differentiating Inh from BL demands appears to be most difficult as the achieved values do not appear to be much above chance level. Statistics revealed though that all achieved accuracies are significantly above chance level, as indicated by the annotations in the table. The influence of CCA as a spatial filter on the classification results in terms of accuracy has also been evaluated in Table 1. An increase in accuracy can be reached in all cases.

**Table 1.** Classification accuracies (and standard deviations) achieved in a 10-fold cross-validation with a linear kernel on 0-1000 ms time frame. One feature set comprised the ERPs of the 17 midline channels, the second feature set comprised these ERPs treated with a CCA as a spatial filter (27x 17). Significances are indicated as follows: * p < .05, ** p < .005, *** p < .0005

| Feature Set |  | Inh vs BL | Up1 vs BL | Up2 vs BL | Up1 vs Inh | Up2 vs Inh |
| --- | --- | --- | --- | --- | --- | --- |
| ERP | Acc | 55.83 % *** | 56.65 % *** | 67.73 % *** | 62.21 % *** | 70.09 % *** |
|  | Stdv | 5.38 | 6.32 | 8.92 | 7.38 | 8.41 |
| ERP (CCA) | Acc | 61.12 % *** | 61.20 % *** | 73.38 % *** | 68.23 % *** | 74.73 % *** |
|  | Stdv | 6.24 | 6.71 | 9.24 | 5.28 | 8.20 |

#### 3.2.2. Classification on pupil diameter

In Table 2 the classification accuracies that can be achieved by using the pupil diameter values only are displayed. Again all values are significantly above chance level. As for the ERP features, differentiating Inh from BL demands appears to be most difficult. For all other comparisons the accuracies that can be achieved with the pupil diameter are even higher than for the ERP features, although the number of used features was considerably smaller.

**Table 2.** Classification results for using the pupil diameter of both eyes as feature set. All displayed results are 10-fold cross validated during SVM classification with a linear kernel. Significances are indicated as follows: * p < .05, ** p < .005, *** p < .0005

| Feature Set |  | Inh vs BL | Up1 vs BL | Up2 vs BL | Inh vs Up1 | Inh vs Up2 |
| --- | --- | --- | --- | --- | --- | --- |
| Pupil | Acc | 53.41 % *** | 61.77 % *** | 77.42 % *** | 60.46 % *** | 75.76 % *** |
|  | Stdv | 5.51 | 8.28 | 9.54 | 8.51 | 11.28 |

#### 3.2.3. Classification on power spectra

Table 3 shows the achieved accuracies when using the power spectra from 4-13 Hz as features in the individual comparisons. The presented accuracies show the same pattern as the ones achieved by using ERP or pupil dilation features, yet they are slightly lower. Nevertheless, all results are statistically significant. In addition to the classification rates, the weights of the used SVM were also evaluated. Figure 5 shows the weights in the interpretable form as suggested by Haufe and colleagues. Figure 5 A provides all values in a heatmap, whereas B and C show the topological distribution of the weights averaged over theta and alpha band power respectively. This analysis reveals that mainly frontal theta and parietal alpha seem to play a role for the distinction. A finer pattern is shown in the topological plots (B) indicating that inhibition seems to be a process that correlates more with central theta synchronization, whereas updating with a little bit more frontal theta activity. The fact that updating and inhibition demands cannot only be distinguished from baseline demands but also from each other can be taken as evidence for the diversity of executive functions with regard to their neurophysiological signatures. The different weights of the used SVMs (power spectra used as features) also provide insights into the feature characteristics of the two executive functions.

**Table 3.**
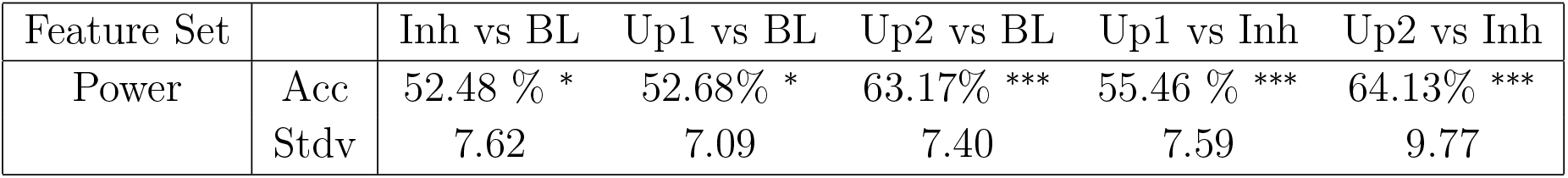
Classification accuracies achieved in a 10-fold cross-validation with a linear kernel on the power spectra between 4-13 Hz, calculated on the 0-1000 ms time frame from stimulus onset. Significances are indicated as follows: * p < .05, ** p < .005, *** p < .0005

| Feature Set |  | Inh vs BL | Up1 vs BL | Up2 vs BL | Up1 vs Inh | Up2 vs Inh |
| --- | --- | --- | --- | --- | --- | --- |
| Power | Acc | 52.48 % * | 52.68% * | 63.17% *** | 55.46 % *** | 64.13% *** |
|  | Stdv | 7.62 | 7.09 | 7.40 | 7.59 | 9.77 |

**Figure 5.**
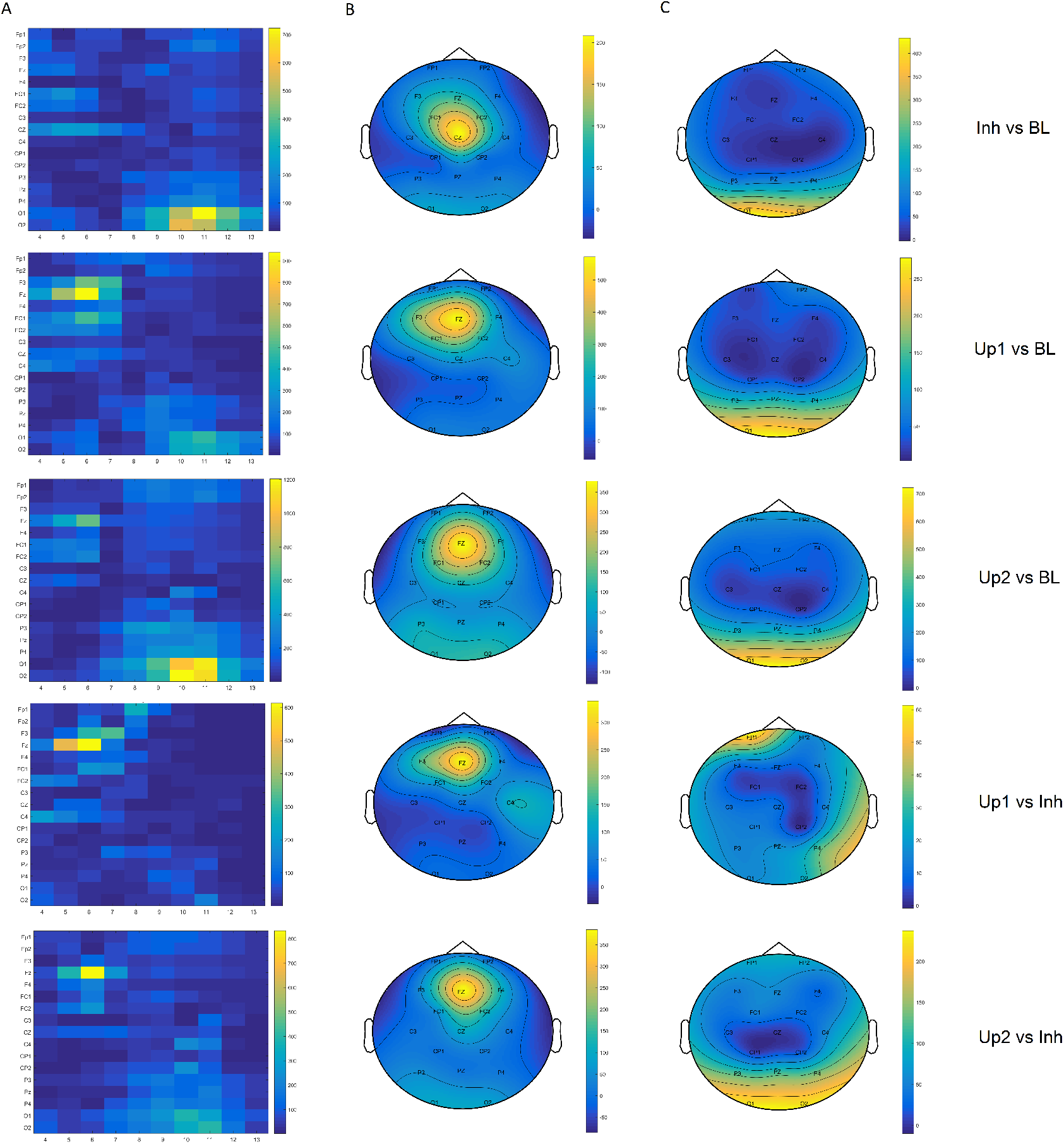
SVM weights in neurophysiological interpretable form, calculated for every subject individually. Shown are weights that are assigned to each feature in the frequency domain according to its importance, averaged over all subjects. A: Displayed are the absolute weights for all features in a heatmap. B: The weights of theta band are averaged over the frequency bins 4 - 7 Hz and displayed in a topological plot. C: Displays the topological distribution of the weights averaged over the alpha band (8 - 13 Hz).

#### 3.2.4. Cross-class classification

Cross-class classifications were performed to reveal if joint feature characteristics in either the spectral or ERP features are shared in large proportions by the two EFs. The thereby obtained results can be seen in Table 4. The reported accuracies describe the percentage of trials of the test condition that have been classified as BL condition. The percentage of trials classified as training condition can be calculated by 100 minus the percentage classified as BL. All results provided are close to random except the Up2 vs BL distinction in the frequency as well as in the time domain. The statistical analysis revealed that the distinction Inh vs Up2 is significantly better than random, whereas all other comparisons are not. The significant effect indicates that a classifier trained on Up2 vs BL trials will classify Inh trials significantly above chance level (around 60 %) as BL trials and not as Up2 trials. So the classifier trained on one EF does not recognize the demands imposed onto another EF. This effect as well as the chance level classifications for the other conditions provides evidence that the neurophysiological signatures of the two EFs are substantially different so that no cross-class classifications are possible.

**Table 4.** The table provides cross-class classification for ERP features as well as for the power spectra. The classifier was trained on EF1 vs BL and tested on trials imposing demands on EF2 only. Therefore, the here presented accuracies represent the percentage of trials classified as BL and 100 - the here displayed percentage reveals the share of trials classified as EF1 respectively. Significances are indicated as follows: * p < .05, ** p < .005, *** p < .0005

| Feature Set | Trainset | Inh vs BL | Inh vs BL | Up1 vs BL | Up2 vs BL |
| --- | --- | --- | --- | --- | --- |
|  | Testset | Up1 | Up2 | Inh | Inh |
| Power | Acc | 51.55 % | 47.86 % | 52.84 % | 61.26 % *** |
|  | Stdv | 9.33 | 12.13 | 8.09 | 9.41 |
| ERP | Acc | 52.93 % | 51.22 % | 49.81 % | 57.92 % *** |
|  | Stdv | 9.37 | 9.33 | 10.45 | 9.56 |

## 4. Discussion

The aim of this paper is to provide neurophysiological evidence for the diversity of executive working memory functions based on the analysis of EEG data. To achieve that standard neurophysiological analysis techniques are combined with machine learning approaches to extract neural correlates of the EFs in EEG data. By this we want to show the benefits of using machine learning in a context like this. It enables to use inter subject variability as an advantage while trying to separate mental states or more general, experimental conditions within EEG data. Mathematical optimization in form of SVMs is applied to the data by taking all available information into account. We aim to establish the usage as a supplementing technique to extract as much information as possible from EEG data, since the methodology provides advantages compared to standard techniques.

### 4.1. Neurophysiological analysis

The standard analysis of the ERPs, averaged over the individual factor levels, reveals an elicited P300 which decreases in amplitude throughout the conditions in the following order: BL > Inh > Up1 > Up2. A similar pattern of results was found in the pupil dilation data as well as in the power spectra. The pupil diameter increases with increasing WML, whereas the power spectra yield well known indicators for WML, namely alpha desynchronization and theta synchronization cf., (Ewing and Fairclough, 2010a; Gevins et al., 1997). The feature sets do not seem to show characteristic patterns which can be used to differentiate the two EFs, even though the comparisons between the conditions revealed statistically significant differences in the ERPs and power spectra at several electrode positions. It is more likely that the differences are due to the general amount of WML that is present throughout the task. Since updating is induced by the main task and inhibition only by a secondary stimulus presentation to which no explicit response was necessary, it can be assumed that the induced load is likely to be higher for updating than for inhibition. Moreover it is uncontroversial that updating demands elicited by the 2-back task are more challenging than the ones elicited by the 1-back task, as more letters need to be constantly updated in WM. A decrease in P300 amplitude, as is present in our data, is also well in line with previous findings from the literature with respect to an increase of overall WML (Ewing and Fairclough, 2010a; Watter et al., 2001). Standard analysis techniques therefore reveal differences between the four factor levels but they can only be linked to the general amount of WML and not to specific properties of the individual EFs.

Despite consistency with literature, another aspect can be considered with respect to the comparably weak correlates of inhibition. In a more recent version of their model Miyake and Friedman (2012) state that the inhibition ability is the core property of all EF and Friedman emphasizes this statement by postulating there is no unique variance describing inhibition (Friedman et al., 2011,0). Potentially, this hypothesis could provide a theoretical explanation why inhibition trials provided the overall weakest results, neurophysiologically as well as in terms of classification accuracy. However, it could also be argued that the secondary nature of the flanker stimulus inducing the inhibition demands is the reason for that. No explicit reaction to this stimulus was demanded in contrast to the central stimulus used for inducing updating demands. Designing a task that puts the inhibition demands more into focus might resolve this issue and reveal clearer and stronger neural correlates for inhibition.

### 4.2. Classification

As a new approach that aims to disentangle individual characteristics of the EFs, we applied machine learning to the data. In this case we chose SVMs, but other methods such as LDA would perform similarly (Lotte et al., 2007). The use of single trial data for machine learning is a necessary step as the ML algorithm needs the single trial data to learn the distribution of the data. One of the benefits of the ML approach compared to standard techniques is, that inter-subject variability is incorporated rather than eliminated from the analysis. Hence differences that show variability across subjects but are present steady within the dataset, will be factored in account when investigating a possible distinction of properties. Therefore, the training for each subject individually, guarantees to find an optimal model for each subject. Another benefit is that a broad number of features from many channels can be taken into account at once. Basis is a mathematical optimization approach, which might be able to reveal patterns that are not visible with conventional methods. With this approach it could be shown that a significantly better than random distinction was possible between the EF conditions on the basis of single trial ERPs and also single trial power spectra. Therefore, in either of the two feature sets differences can be found that enable a distinction between the two EFs and between EFs and baseline demands. The fact that updating and inhibition demands cannot only be distinguished from baseline demands but also from each other speaks in favor of a diversity of executive functions with regard to their neurophysiological signatures. The achieved accuracies measure the success of the distinction, which is higher for single trial ERPs and mean pupil diameter than for the power spectra. A general observation that can be made is that the accuracy values mirror the gradient that was found in the measured physiological signals. The bigger the difference in induced WML between the conditions, the higher the accuracy for the respective distinction by SVM classification for all three feature sets. To rule out that the differences between the conditions rely on the amount of induced WML only and not on individual variance caused by the EFs, a cross-class classification was performed. Cross-class classification indicates how much the characteristics of the EFs overlap. If signal strength (reflected by ERP amplitude, ERD/ERS) would be the only difference between the conditions, the overlap should be large and cross-class classification accuracies should be significantly above chance level. Cross-class classification was tested for ERP and spectral features and in both cases three out of four tests provided accuracies around chance level. The only exception was a classifier trained on Up2 vs BL that was tested on Inh trials. It turned out that Inh is rather classified as BL than as Up2, revealing that Inh seems to be closer to BL demands than to Up2 demands. This effect and the fact that in all other cases only random accuracies could be achieved in cross-class classification provides evidence for the hypothesis that there are larger differences in the signals reflecting different EFs than those visible when using conventional methods for neurophysiological analysis.

In addition to this, the weights of the SVM classification approach (based on power spectra features) were inspected more closely to find out which features are prominently used in the distinction. The resulting values indicate that especially features in the occipital/parietal alpha and in the frontal theta yielded the highest weights. These are also features that are known from the literature to strongly correlate with WML. Apart from these WML related features, no other features seem to play a prominent role according to the interpretable weights (see Figure 5). Thus, it seems that the SVM classification for the analysis of the diversity of the two EFs takes only the relevant neural signatures related to WML into account. The inspection of the SVM weights also revealed different pattern that shows differences in the theta band power for updating and inhibition. Inhibition correlates with central theta, whereas updating with a more frontal theta band power synchronization. This difference in feature characteristics renders the two EFs differentiable on the basis of their neural signatures, thereby, accounting for the diversity of the two EFs. Yet, the same values reveal a common correlation with occipital/paretial alpha desynchronization in both EF, hence accounting for the unity of these functions. Since the experimental design was chosen very carefully reducing all non-EF related variance to a minimum, it seems legitimate to assume that the discovered differences in the patterns can be traced back to the EFs and not to any confounds from external stimuli. Overall, we conclude that both aspects of the theoretical model put forward by Miyake can be confirmed with actual brain data. We also conclude that using machine learning in addition to classical analysis approaches is a valuable and maybe even necessary extension when aiming to answer psychological questions as has already been promoted by Yarkoni and Westfall (2017).

### 4.3. Limitations

Although it could be demonstrated, that both individual EFs were characterized by their own variance separating it from the other, no claim can be made yet with regard to shifting demands, to fully prove or disprove the statements by Miyake and Friedman. Only a rather strong conclusion can be drawn, with regard to updating and inhibition, namely, that the two executive functions are implemented by two different processes that can be distinguished from each other by means of their underlying neurophysiological signatures. To further confirm the model of Miyake and colleagues, the third executive function shifting also needs to be evaluated. This requires to design a new task, integrating all three executive functions in one experimental setup, to ensure equal conditions for all comparisons and distinctions. It is necessary to keep non-EF variance at a minimum to avoid confounds and to ensure the validity of the analysis. Based on this setup it can be analyzed whether shifting demands also have distinct neural signatures that can be separated from inhibition and updating demands to further demonstrate the diversity aspect of EFs. Analyzing shifting demands might also shed new light on the unity aspect of EFs that could be demonstrated in the current study. In addition to this it might also be useful to design similar experiments in which the EFs are induced by different task. Despite being known for inducing load on a specific EF, it can be assumed that non of the tasks is of such a singular nature, that only one resource is used. Applying this approach on a task battery will reveal a better picture with respect to the diversity of EFs.

A general remark is that the presented approach can only be seen as a supplementing method, but not as a standalone technique. It only makes sense to apply ML to data when it has been ensured that existing variance in the data is only due to the mental states of interest and not to artifacts introduced by visual presentation or other artifacts of any kind. To this end, it also needs to be acknowledged that classification on pupil data was done to see how well this simple feature correlates with the WML. It is clear that the pupil diameter cannot be a indicator for EFs but only for WML in general. Still, the high correlation with the overall load and the high classification accuracies when using the pupil diameter as a feature, opens up the possibility to use it as a simple measure for WML detection or as an additional measure to EEG to stabilize or even improve WML detection on the basis physiological features.

## 5. Conclusion

The approach chosen in this paper to disentangle neurophysiological signatures of cognitive functions by means of machine learning provide insights that support the theoretical model of Miyake and colleagues describing the unity and diversity of EFs. It can be shown that the two executive functions updating and inhibition, which both induce WML, can be separated on the basis of single trial ERPs and power spectra. Using power spectra yielded less accurate results but allowed to apply a new method developed by Haufe and colleagues to reveal patterns in the spectra that can be extracted and linked to the two individual executive functions. Inhibition is characterized by an increased frontal activity in the theta band, whereas updating demands are characterized by an increased central activity in the theta band. The results therefore, substantiate the hypothesis that the two executive functions should be considered as two separable processes in WM. Our approach of applying machine learning techniques to neurophysiological data in order to substantiate theoretical distinctions in functional models can also be a promising approach for other research areas in cognitive neuroscience. It supplements the classical approaches and is thereby able to extract more knowledge out of existing data, then standard analysis techniques can provide.

## Acknowledgments

This study was funded by the Leibnitz Science Campus Tübingen ‘Informational Environments’. Tanja Krumpe is a doctoral student at the LEAD Graduate School & Research Network [GSC1028], funded by the Excellence Initiative of the German federal and state governments.

## References

Allison, B. Z. and Polich, J. (2008). Workload assessment of computer gaming using a single-stimulus event-related potential paradigm. Biological psychology, 77(3):277–283.

Baddeley, A. (1996). Exploring the central executive. The Quarterly Journal of Experimental Psychology: Section A, 49(1):5–28.

Baddeley, A. (2007). Working memory, thought, and action. OUP Oxford.

Baddeley, A. (2012). Working memory: theories, models, and controversies. Annual review of psychology, 63:1–29.

Baddeley, A., Hitch, G., et al. (1974). Working memory. The psychology of learning and motivation, 8:47–89.

Barrouillet, P., Bernardin, S., and Camos, V. (2004). Time constraints and resource sharing in adults’ working memory spans. Journal of Experimental Psychology: General, 133(1):83.

Beatty, J. and Lucero-Wagoner, B. (2000). The pupillary system. Handbook of psychophysiology, 2:142–162.

Botvinick, M. M., Braver, T. S., Barch, D. M., Carter, C. S., and Cohen, J. D. (2001). Conflict monitoring and cognitive control. Psychological review, 108(3):624.

Brouwer, A.-M., Hogervorst, M. A., Van Erp, J. B., Heffelaar, T., Zimmerman, P. H., and Oostenveld, R. (2012). Estimating workload using eeg spectral power and erps in the n-back task. Journal of neural engineering, 9(4):045008.

Burgess, P. W. (1997). Theory and methodology in executive function research. Methodology of frontal and executive function, pages 81–116.

Chang, C.-C. and Lin, C.-J. (2011). LIBSVM: A library for support vector machines. ACM Transactions on Intelligent Systems and Technology, 2:27:1–27:27. Software available at http://www.csie.ntu.edu.tw/cjlin/libsvm.

Chatham, C. H., Herd, S. A., Brant, A. M., Hazy, T. E., Miyake, A., O’Reilly, R., and Friedman, N. P. (2011). From an executive network to executive control: a computational model of the n-back task. Journal of cognitive neuroscience, 23(11):3598–3619.

Chen, Y., Mitra, S., and Schlaghecken, F. (2007). Interference from the irrelevant domain in n-back tasks: an erp study. Acta Neurologica Taiwanica, 16(3):125.

Chen, Y., Mitra, S., and Schlaghecken, F. (2008). Sub-processes of working memory in the n-back task: An investigation using erps. Clinical Neurophysiology, 119(7):1546–1559.

Collette, F., Hogge, M., Salmon, E., and Van der Linden, M. (2006). Exploration of the neural substrates of executive functioning by functional neuroimaging. Neuroscience, 139(1):209–221.

Collette, F., Van der Linden, M., Laureys, S., Delfiore, G., Degueldre, C., Luxen, A., and Salmon, E. (2005). Exploring the unity and diversity of the neural substrates of executive functioning. Human brain mapping, 25(4):409–423.

Cowan, N., Elliott, E. M., Saults, J. S., Morey, C. C., Mattox, S., Hismjatullina, A., and Conway, A. R. (2005). On the capacity of attention: Its estimation and its role in working memory and cognitive aptitudes. Cognitive psychology, 51(1):42–100.

Cowan, N., Saults, J. S., and Blume, C. L. (2014). Central and peripheral components of working memory storage. Journal of Experimental Psychology: General, 143(5):1806.

Davelaar, E. J. (2012). When the ignored gets bound: sequential effects in the flanker task. Frontiers in psychology, 3.

Diamond, A. (2013). Executive functions. Annual review of psychology, 64:135.

Ecker, U. K., Lewandowsky, S., Oberauer, K., and Chee, A. E. (2010). The components of working memory updating: An experimental decomposition and individual differences. Journal of Experimental Psychology: Learning, Memory, and Cognition, 36(1):170.

Eriksen, C. W. (1995). The flankers task and response competition: A useful tool for investigating a variety of cognitive problems. Visual Cognition, 2(2–3):101–118.

Ewing, K. and Fairclough, S. (2010a). The impact of working memory load on psychophysiological measures of mental effort and motivational disposition. Human Factors: A system view of human, technology and organisation. Maastricht: Shaker Publishing.

Ewing, K. C. and Fairclough, S. H. (2010b). The effect of an extrinsic incentive on psychophysiological measures of mental effort and motivational disposition when task demand is varied. In Proceedings of the human factors and ergonomics society annual meeting, volume 54, pages 259–263. Sage Publications Sage CA: Los Angeles, CA.

Friedman, N. P., Miyake, A., Robinson, J. L., and Hewitt, J. K. (2011). Developmental trajectories in toddlers’ self-restraint predict individual differences in executive functions 14 years later: a behavioral genetic analysis. Developmental psychology, 47(5):1410.

Friedman, N. P., Miyake, A., Young, S. E., DeFries, J. C., Corley, R. P., and Hewitt, J. K. (2008). Individual differences in executive functions are almost entirely genetic in origin. Journal of Experimental Psychology: General, 137(2):201.

Gevins, A. and Smith, M. E. (2000). Neurophysiological measures of working memory and individual differences in cognitive ability and cognitive style. Cerebral cortex, 10(9):829–839.

Gevins, A., Smith, M. E., McEvoy, L., and Yu, D. (1997). High-resolution eeg mapping of cortical activation related to working memory: effects of task difficulty, type of processing, and practice. Cerebral cortex, 7(4):374–385.

Good, P. (2013). Permutation tests: a practical guide to resampling methods for testing hypotheses. Springer Science & Business Media.

Gratton, G., Coles, M. G., and Donchin, E. (1992). Optimizing the use of information: strategic control of activation of responses. Journal of Experimental Psychology: General, 121(4):480.

Haufe, S., Meinecke, F., Görgen, K., Dähne, S., Haynes, J.-D., Blankertz, B., and Bießmann, F. (2014). On the interpretation of weight vectors of linear models in multivariate neuroimaging. Neuroimage, 87:96–110.

Jasper, H. (1958). The 10/20 international electrode system. EEG and Clinical Neurophysiology, 10:371–375.

Jensen, O. and Tesche, C. D. (2002). Frontal theta activity in humans increases with memory load in a working memory task. European journal of Neuroscience, 15(8):1395–1399.

Jonides, J., Schumacher, E. H., Smith, E. E., Lauber, E. J., Awh, E., Minoshima, S., and Koeppe, R. A. (1997). Verbal working memory load affects regional brain activation as measured by pet. Journal of cognitive neuroscience, 9(4):462–475.

Kane, M. J., Hambrick, D. Z., Tuholski, S. W., Wilhelm, O., Payne, T. W., and Engle, R. W. (2004). The generality of working memory capacity: a latent-variable approach to verbal and visuospatial memory span and reasoning. Journal of Experimental Psychology: General, 133(2):189.

Karatekin, C., Marcus, D. J., and Couperus, J. W. (2007). Regulation of cognitive resources during sustained attention and working memory in 10-year-olds and adults. Psychophysiology, 44(1):128–144.

Krause, C. M., Pesonen, M., and Hämäläinen, H. (2010). Brain oscillatory 4–30 hz electroencephalogram responses in adolescents during a visual memory task. Neuroreport, 21(11):767–771.

Krause, J., Taylor, J. G., Schmidt, D., Hautzel, H., Mottaghy, F. M., and Müller-Gärtner, H.-W. (2000). Imaging and neural modelling in episodic and working memory processes. Neural Networks, 13(8):847–859.

Laeng, B., Ørbo, M., Holmlund, T., and Miozzo, M. (2011). Pupillary stroop effects. Cognitive processing, 12(1):13–21.

Lotte, F., Congedo, M., Lécuyer, A., Lamarche, F., and Arnaldi, B. (2007). A review of classification algorithms for eeg-based brain–computer interfaces. Journal of neural engineering, 4(2):R1.

MATLAB (2015). version 7.10.0 (R2015b). The MathWorks Inc., Natick, Massachusetts.

Missonnier, P., Deiber, M.-P., Gold, G., Millet, P., Pun, M. G.-F., Fazio-Costa, L., Giannakopoulos, P., and Ibáñez, V. (2006). Frontal theta event-related synchronization: comparison of directed attention and working memory load effects. Journal of Neural Transmission, 113(10):1477–1486.

Miyake, A. and Friedman, N. P. (2012). The nature and organization of individual differences in executive functions four general conclusions. Current directions in psychological science, 21(1):8–14.

Miyake, A., Friedman, N. P., Emerson, M. J., Witzki, A. H., Howerter, A., and Wager, T. D. (2000). The unity and diversity of executive functions and their contributions to complex frontal lobe tasks: A latent variable analysis. Cognitive psychology, 41(1):49–100.

Miyake, A. and Shah, P. (1999). Models of working memory: Mechanisms of active maintenance and executive control. Cambridge University Press.

Monsell, S. (2003). Task switching. Trends in cognitive sciences, 7(3):134–140.

Nichols, T. E. and Holmes, A. P. (2002). Nonparametric permutation tests for functional neuroimaging: a primer with examples. Human brain mapping, 15(1):1–25.

Oberauer, K. (2009). Design for a working memory. Psychology of learning and motivation, 51:45–100.

Owen, A. M., McMillan, K. M., Laird, A. R., and Bullmore, E. (2005). N-back working memory paradigm: A meta-analysis of normative functional neuroimaging studies. Human brain mapping, 25(1):46–59.

Pesonen, M., Hämäläinen, H., and Krause, C. M. (2007). Brain oscillatory 4–30 hz responses during a visual n-back memory task with varying memory load. Brain research, 1138:171–177.

Platt, J. C. (2000). Probabilistic Outputs for Support Vector Machines and Comparisons to Regularized Likelihood Methods. MIT Press.

Pratt, N., Willoughby, A., and Swick, D. (2011). Effects of working memory load on visual selective attention: behavioral and electrophysiological evidence. Frontiers in human neuroscience, 5:57.

Putze, F., Jarvis, J.-P., and Schultz, T. (2010). Multimodal recognition of cognitive workload for multitasking in the car. In Pattern Recognition (ICPR), 2010 20th International Conference on, pages 3748–3751. IEEE.

Sanders, A. and Lamers, J. (2002). The eriksen flanker effect revisited. Acta Psychologica, 109(1):41–56.

Scharinger, C., Soutschek, A., Schubert, T., and Gerjets, P. (2015). When flanker meets the n-back: What eeg and pupil dilation data reveal about the interplay between the two central-executive working memory functions inhibition and updating. Psychophysiology, 52(10):1293–1304.

Shallice, T. and Burgess, P. (1991). Deficits in strategy application following frontal lobe damage in man. Brain, 114(2):727–741.

Shallice, T. and Burgess, P. (1993). Supervisory control of action and thought selection.

Spüler, M., Walter, A., Rosenstiel, W., and Bogdan, M. (2014). Spatial filtering based on canonical correlation analysis for classification of evoked or event-related potentials in eeg data. IEEE Transactions on Neural Systems and Rehabilitation Engineering, 22(6):1097–1103.

Stipacek, A., Grabner, R., Neuper, C., Fink, A., and Neubauer, A. (2003). Sensitivity of human eeg alpha band desynchronization to different working memory components and increasing levels of memory load. Neuroscience Letters, 353(3):193–196.

Unsworth, N. and Engle, R. W. (2007). The nature of individual differences in working memory capacity: active maintenance in primary memory and controlled search from secondary memory. Psychological review, 114(1):104.

Vapnik, V. and Chervonenkis, A. (1974). Theory of Pattern Recognition. Technical report.

Walter, C., Rosenstiel, W., Bogdan, M., Gerjets, P., and Spüler, M. (2017). Online eeg-based workload adaptation of an arithmetic learning environment. Frontiers in Human Neuroscience, 11:286.

Watter, S., Geffen, G. M., and Geffen, L. B. (2001). The n-back as a dual-task: P300 morphology under divided attention. Psychophysiology, 38(6):998–1003.

Yarkoni, T. and Westfall, J. (2017). Choosing prediction over explanation in psychology: Lessons from machine learning. Perspectives on Psychological Science, 12(6):1100–1122.

